# Dynamic yield responses of chickpea (*Cicer arietinum*) to terminal drought are accompanied by changes in grain composition

**DOI:** 10.64898/2026.02.26.708401

**Authors:** Paul Hopgood, Sally Buck, Melissa Bain

## Abstract

Chickpea is predominantly grown under rainfed conditions in regions where terminal drought limits yield, yet little is known about how this stress influences both vegetative allocation and reproductive dynamics leading to altered grain composition. We imposed a controlled terminal drought, with a rewatered treatment group, on three Desi cultivars (ICC4958, ICC1882 and CBA Captain) reported to have contrasting drought tolerance, quantifying vegetative biomass, reproductive node productivity across developmental regions and grain macronutrient composition. Under drought, vegetative responses reflected genotype-specific resource partitioning strategies particularly evident in severe root degradation and increase stem dry matter content that was only partially alleviated in rewatered plants. Reproductive outcomes were strongly influenced by developmental stage at the time of stress, with increased pod abortion observed particularly at nodes initiating seed development under drought treatment. Grain composition of seeds filled under drought was significantly altered by stress, with increased protein concentration and decreased starch content under both Drought and Recovery treatments independent of cultivar, likely due to water limitation at crucial filling stages. These findings demonstrate that the developmental timing of terminal drought interacts with cultivar growth strategy to influence pod production and grain nutritional quality in chickpea.

**Highlight:** The developmental timing of terminal drought interacts with cultivar-dependent growth strategies to influence pod productivity and grain nutritional quality in chickpea.

## Introduction

Chickpea (*Cicer arietinum*) has been an important legume in traditional food systems for thousands of years and continues to have increasing significance to sustainable global agriculture, food security and nutrition (Bar-El Dadon et al., 2017; Copperstone et al., 2025). Originating in the Mediterranean, chickpeas are widely cultivated in predominantly arid and semi-arid regions of the Middle East, North and East Africa, Central and South Asia, as well as sub-tropical and more temperate regions of Australia (Igolkina et al., 2023; Varshney et al., 2019). Whilst the nutritional value of chickpea has been understood by many food cultures, our knowledge of the mechanisms by which the complement of nutrients and bio-actives found in these grains contribute to human health is increasing rapidly (Begum et al., 2023; Copperstone et al., 2025; Elma Mathew & Shakappa, 2022; Wang et al., 2021). The combination of dietary fibres, bioactive peptides and other compounds including flavonoids found in chickpea have been associated with improved gut health, acting to mitigate risks of chronic cardiovascular disease and some cancers (Begum et al., 2023; Copperstone et al., 2025; Elma Mathew & Shakappa, 2022; Wang et al., 2021). Given the importance of these grain components to the provision of basic nutrition as well as in human health, understanding the impacts of environment on grain development and composition will be key to securing their nutritional value. In the climates in which chickpea is grown rain fed systems dominate and crops are often challenged by terminal drought which has significant impacts on production and has become more urgent challenge in a changing climate (Arriagada et al., 2022; Farooq et al., 2017; Jha et al., 2020; Karalija et al., 2022). One cultivar which has become important in breeding for and understanding of terminal drought resistance is ICC4958 (International Crops Research Institute for the Semi-Arid Tropics (ICRISAT), 1992), which although only moderately resistant in large germplasm screens (Krishnamurthy et al., 2010) has well characterised physiology under stress. Drought resistance in ICC4958 has been attributed largely to its ability to produce greater root length density (RLD) and deeper root systems under terminal drought stress than other cultivars (Kashiwagi et al., 2005, 2006). In the field this root architecture, in cultivars including ICC4958, translates to increased soil water uptake from layers deeper than 45cm under stress, which in the context of other water use responses in the canopy contributes to their drought resistance (Purushothaman et al., 2016; Ramamoorthy et al., 2017). Growth patterns in shoots also influence performance under drought with conservative investment in leaf area during vegetative growth and the ability to support greater shoot biomass during reproduction central to drought resistance in chickpea (Ramamoorthy et al., 2016). ICC1882 is a drought susceptible variety, and has been used as a parent in mapping populations with ICC4958 (Varshney et al., 2014) which have identified quantitative trait loci (QTL) connected to root traits (Jaganathan et al., 2015; Singh et al., 2016), canopy size and conductance characteristics (Sivasakthi et al., 2018), as well as 100 seed weight (Kale et al., 2015). This comparison between ICC4958 and ICC1882 has also generated an expressed sequence tag (EST) collection (Varshney et al., 2009) and transcriptome maps across development (Garg et al., 2016) for drought responsive genes.

This study aims to characterise impacts of an acute, terminal drought on the productivity of reproductive nodes and quality of grain produced in the benchmark ICC4958 and ICC1882 cultivars. To compare these effects in germplasm bred for a different growing region, we also include an elite cultivar Chickpea Breeding Australia (CBA) Captain which has broad adaptation for the sub-tropical region of Northern Australia, and phenology that is early to mid-maturing (Hobson et al., 2021). There are few studies in chickpea which consider protein content (Benali et al., 2023; Samineni et al., 2022; Varol et al., 2020), starch (Awasthi et al., 2014; Hossein Behboudian et al., 2001) or comprehensive proximal composition (Awasthi et al., 2024; Nayyar et al., 2006) under drought stress, and to our knowledge none which have investigated this in ICC4958 and ICC1882 despite extensive physiological characterisation. There is however increasing evidence that resilience in carbohydrate metabolism is important to drought tolerance mechanisms in chickpea (Farooq et al., 2018; Kudapa et al., 2024; Singh et al., 2023) and that this has impacts on grain filling (Awasthi et al., 2014, 2024; Nayyar et al., 2006). We investigate the impacts of terminal drought and rewatering on the composition of grain filled under stress, comparing the drought tolerant, susceptible and broadly adapted cultivars. Analysis of grain composition is placed in the context of temporal effects of drought stress on reproduction, by separation of physiological measurements along the branches corresponding to key stages of treatments and development.

## Methods

### Cultivars

Three Desi chickpeas were selected for this experiment; ICC4958 a drought resistant accession which is high yielding under terminal drought (International Crops Research Institute for the Semi-Arid Tropics (ICRISAT), 1992); a drought susceptible accession ICC1882; and CBA Captain, a key variety in Northern Australian regions bred for broad environmental adaptation (Hobson et al., 2021). Seed was sourced from materials held at the Commonwealth Scientific and Industrial Research Organisation (CSIRO) (Buck et al., 2025), with the exception of CBA Captain which was obtained from Delta Agribusiness (https://www.deltaag.com.au/).

### General growth conditions

Seeds were planted at 2cm depth into a 50:50 (w/w) mix of compost and sand in free-draining 23cm pots, and inoculated with EasyRhiz N type inoculum (New Edge Microbials) containing *Mesorhizobium ciceri* strain CC1192. So that no nutrient would be limiting soil was also supplemented with controlled release fertiliser (Osmocote, Stotts). Plants were grown in glasshouses from November to January in Canberra, Australia at a daytime average temperature of 22°C and nighttime average of 15 °C. Plants were sprinkler watered every second day up until treatments were applied.

### Biomass measurements

Peak biomass measurements were taken for each cultivar on plants grown alongside those used for the drought trial described below (n=3), under general growth conditions in randomised blocks, and destructively harvested a week before the trial commenced. Total leaf area per plant was measured using a Licor leaf area meter (LI-3100C). To calculate relative water content (RWC) three fully developed leaves were selected at the tip of a branch and weighed fresh (FW), then submerged overnight in deionised water to obtain turgid weight (TW), before drying for 5 days at 37C to acquire dry weights (DW) (n=3). The leaf RWC was calculated as RWC = (FW-DW)/(TW-DW).

For both peak biomass and harvest biomass the total stem and roots, which had been washed in water to remove soil, were weighed fresh at the time of harvest then dried at 37°C for 5 days before DW was obtained. The dry matter content (DMC) of each tissue type was calculated as DMC = DW/FW.

### Terminal drought experiment

Plants were maintained under the general growth conditions described above until the treatments were initiated for all cultivars at 10 days after CBA Captain replicates had entered flowering. Plants were arranged in randomised blocks and a total of 5 or 6 biological replicates sewn for each treatment and cultivar combination.

Due to the indeterminate nature of flowering, tags were used to demark the temporal and developmental regions of each reproductive stem at key stages of treatments. Before the initiation of the treatments black tags were added to all stems with reproductive nodes on the internode above the last formed pod or flower to demark Region 1 (R1) in which pods would fill under treatment conditions (**Figure 1**).

**Figure 1.**
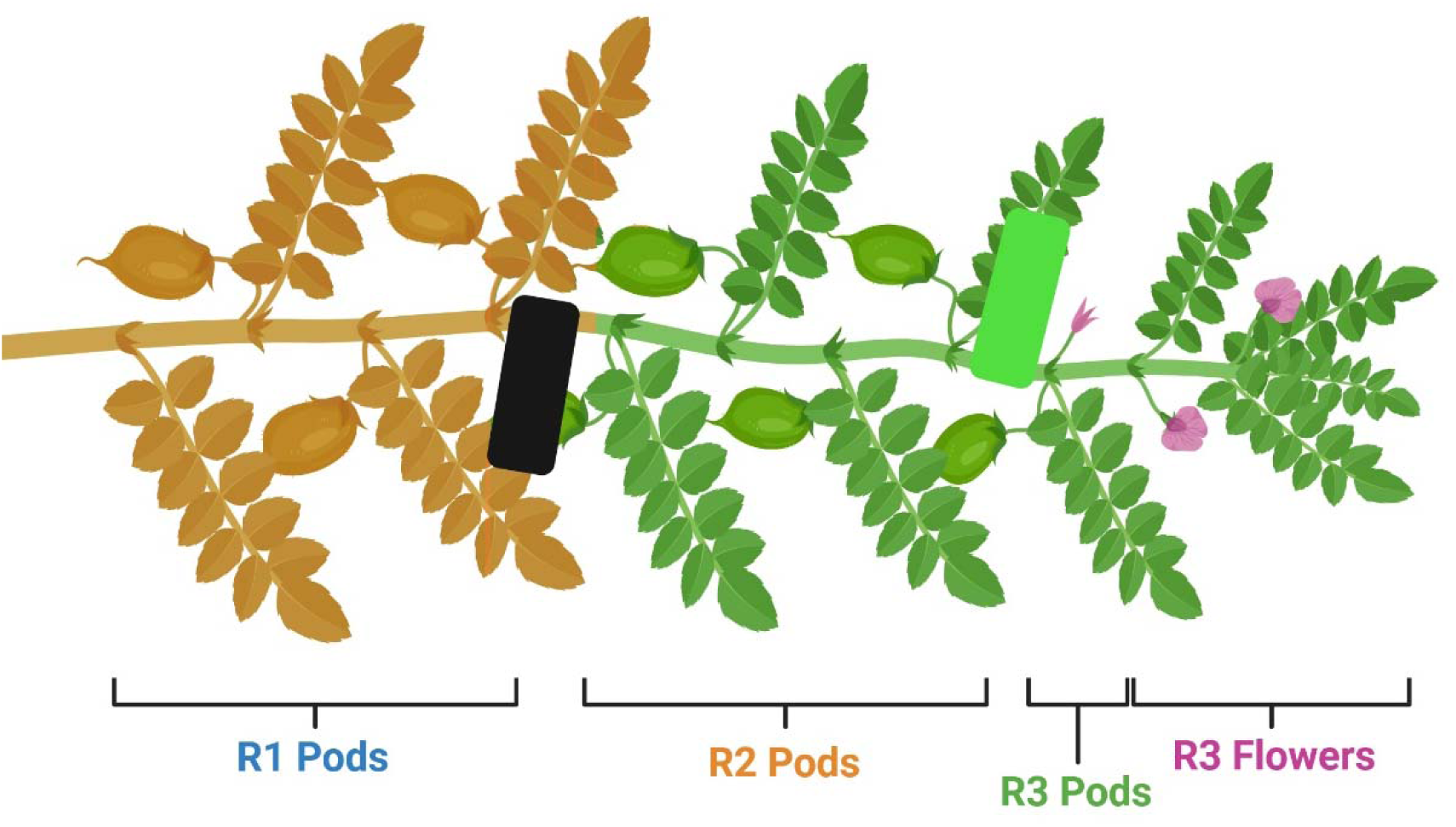
Schematic showing a chickpea stem at the end of flowering to indicate Regions (R1-3) sampled in the drought experiment. The earliest pods to form are shown in senescence (light brown) whilst a range of filling pods and new flowers are present at later stages (green). Tags were applied at key events in the treatments to indicate both temporal and developmental stages of reproductive nodes. A black tag was applied at the initiation of drought treatment, indicating R1 pods set prior to and filled under treatment. The R2 represents pods set under treatment up to rewatering which was marked with a green tag. All pods and flowers set after the green tag are R3, representing reproductive nodes active after either rewatering or continued drought. Created in https://BioRender.com.

Control plants (n=5) were supplied with water equal to the average daily use across cultivars in the 3-4 days before initiation of drought treatments, calculated as the difference between field capacity at saturation (g) and weight after 24hrs before subsequent watering (g). Drought conditions were applied to plants by gradual loss of soil moisture from the induction of treatment to harvest following the method described by Deokar et al. (2011) to produce discrete terminal drought stress suitable for molecular analysis. The rate of water loss under drought conditions was controlled by a reduction of supplied water of 80g subtracted from the average daily water use (Deokar et al., 2011), which was monitored in a random subset of replicates (n=3) by treatment group.

After 10 days of treatment half of the Drought group were rewatered to field capacity then supplied with to the same water as Controls for the remainder of the experiment, and designated Recovery thereafter (n=5). Treatment continued with no change for the remaining Drought group (n=6). At this point green tags were applied to all reproductive stems to indicate the transition between R2 which had set pods in the initial 10 days of treatment and subsequent nodes in R3 which would set flowers and pods under either the rewatered or continued drought conditions (**Figure 1**).

The experiment concluded once CBA Captain Control plants had entered reproductive senescence, at 32 days after initiation of drought treatments.

### Harvest

Seeds were harvested by hand, and pods removed before snap freezing in liquid nitrogen for storage at −80°C until processing. Pod and seed number as well as the number and type of reproductive node were also recorded by developmental region.

### Composition analysis

Seeds were freeze dried whole and weighed by Region for calculation of 100 seed weight (g/100). R1 seed was ground in a Qiagen TissueLyzer II using a 2cm ball bearing at 25Hz for at least 2mins (n=3). The composition of flour was then analysed for total protein, total starch, 80% ethanol soluble and insoluble carbohydrates and total lipid following the procedures described in Buck et al. (2025). In brief, 10mg (+/− 0.5mg) of flour was double extracted by steeping in 50mM Tris-HCl (pH8.0) and 85mM NaCl for 20mins at 25°C to solubilise protein and cleared at 16,000xg before quantitation using Bradford assay (BioRad #5000006). Carbohydrates were sequentially extracted beginning with three washes with 80% ethanol at 32°C for 5mins and centrifugation at 16,000xg. The 80% soluble carbohydrate was quantified from the pooled supernatant using anthrone as in Zhang et al. (2021). Insoluble pellets were analysed using the Megazyme Total Starch kit (#K-TSTA), to enzymatically extract and quantify starch-derived glucose from the pellet. Remaining insoluble material was treated with 1mg Proteinase K (Worthington PROKR, >20U/mg) at 37°C overnight, digested protein removed at 16,000xg and pellet freeze dried for weighing. Total lipids were extracted from 50mg (+/− 0.5mg) using chloroform:methanol at 2:1 (v/v) and phase separation with 150mM ammonium acetate, followed by a second extraction in chloroform only. Lipid extracts were dried under nitrogen and weighed.

### Data analysis

Statistical analyses were performed in RStudio 4.5.1 (Posit team, 2025), in general using linear models with analysis of variance (ANOVA) and Tukey’s post-hoc comparisons where appropriate. For harvest root data with interactions found in ANOVA the simple main effects were analysed with aide from the emmeans package (Lenth, 2025). The analyses deployed for yield data comprised a combination of linear mixed models (LMM) created using the lme4 package (Bates et al., 2015, 2025) for quantitative yield measures, and generalised linear mixed models (GLMM) with negative binomial distribution using the glmmTMB package (Brooks et al., 2025, 2017; McGillycuddy et al., 2025) for discreate yield measures, both of which included random effects to account for the resampling of individual pots across Regions. Finally, multivariate ANOVA (MANOVA) was used to analyse the grain composition data to account for the likely interrelated effects of each component on the total flour from which they were sampled.

## Results

### Cultivar differences affect allocation of peak biomass

Several measures at peak biomass were used to assess how cultivars were allocating biomass under the growth conditions used immediately prior to initiation of treatment. At peak biomass the drought resistant ICC4958 produced a canopy with a significantly smaller total leaf area (mm^2^) (p=0.019) (**Figure 2A**) than both other cultivars, but the RWC of leaves was not found to vary significantly (p>0.05) between the cultivars (**Figure 2B**). CBA Captain plants were larger on average with significantly higher total root dry matter than both ICC4958 and ICC1882 (p=0.010) (**Figure 2D**), and the highest mean stem dry matter, although this difference was not found to be statistically significant between cultivars (p>0.05) (**Figure 2C**)). For all cultivars the proportion of dry matter content in stem and roots was similar at peak biomass, although more variable for both measures in ICC4958 (**Figure 2E,F**).

**Figure 2.**
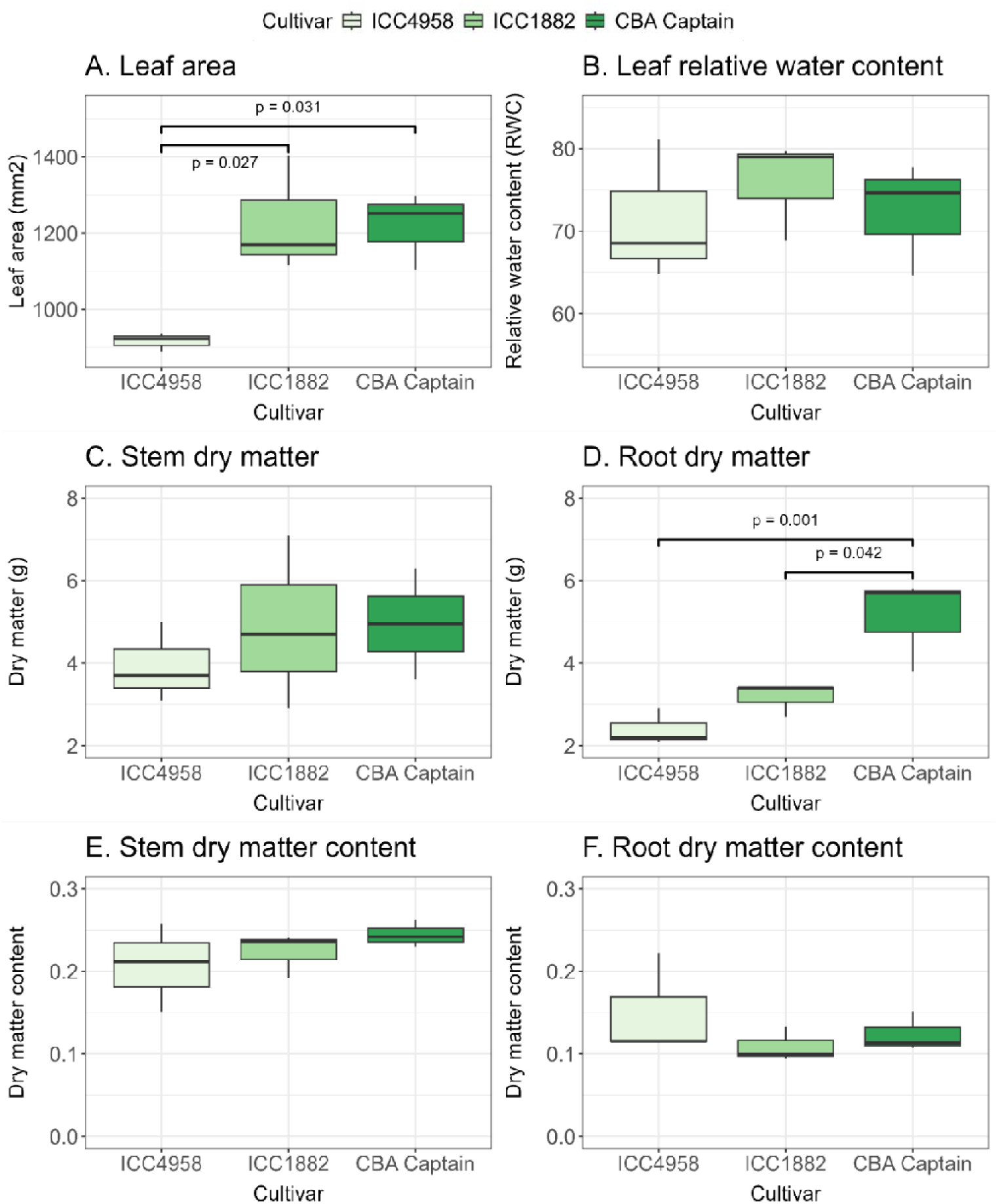
Comparison of peak biomass accumulation across cultivars when grown under control conditions. Total leaf area per plant (**A**) was measured using a Leaf areameter (LI-3100C) on fresh tissue immediately after harvest (n=3). Relative water content of leaves (**B**) was calculated proportionate to turgid weight (w/w) (n=3). The dry weights of both total stems per plant (**C**) and root matter per plant (**D**) were obtained as estimates of aerial and below ground biomass (n=3). In each panel the brackets above the boxplots indicate results of single factor ANOVA with Tukey’s post-hoc comparisons with p-values indicated where significant (α=0.05).

### Harvest biomass indicates cultivar responses to drought stress vary

Differences in final biomass at harvest were observed between cultivars, with treatment effects most evident in the dry matter content of stems and the condition of root biomass. At harvest the stem biomass of ICC1882 was found to be significantly lower than both ICC4958 (p<0.001) and CBA Captain (p<0.01), whilst the effect of treatment was marginal to significant for Drought compared with Control (p=0.058) across all cultivars (**Figure 3A**). The effect of treatment was evident however in the dry matter content (DMC) of stems at harvest, which was significantly higher in Drought than Control (p<0.001) and Recovery (p<0.001) across all cultivars consistent with the protracted desiccation of this group compared with the more moderate conditions for rewatered plants (**Figure 3C**), and evident upon visual inspection of plants (**Figure 4**).

**Figure 3.**
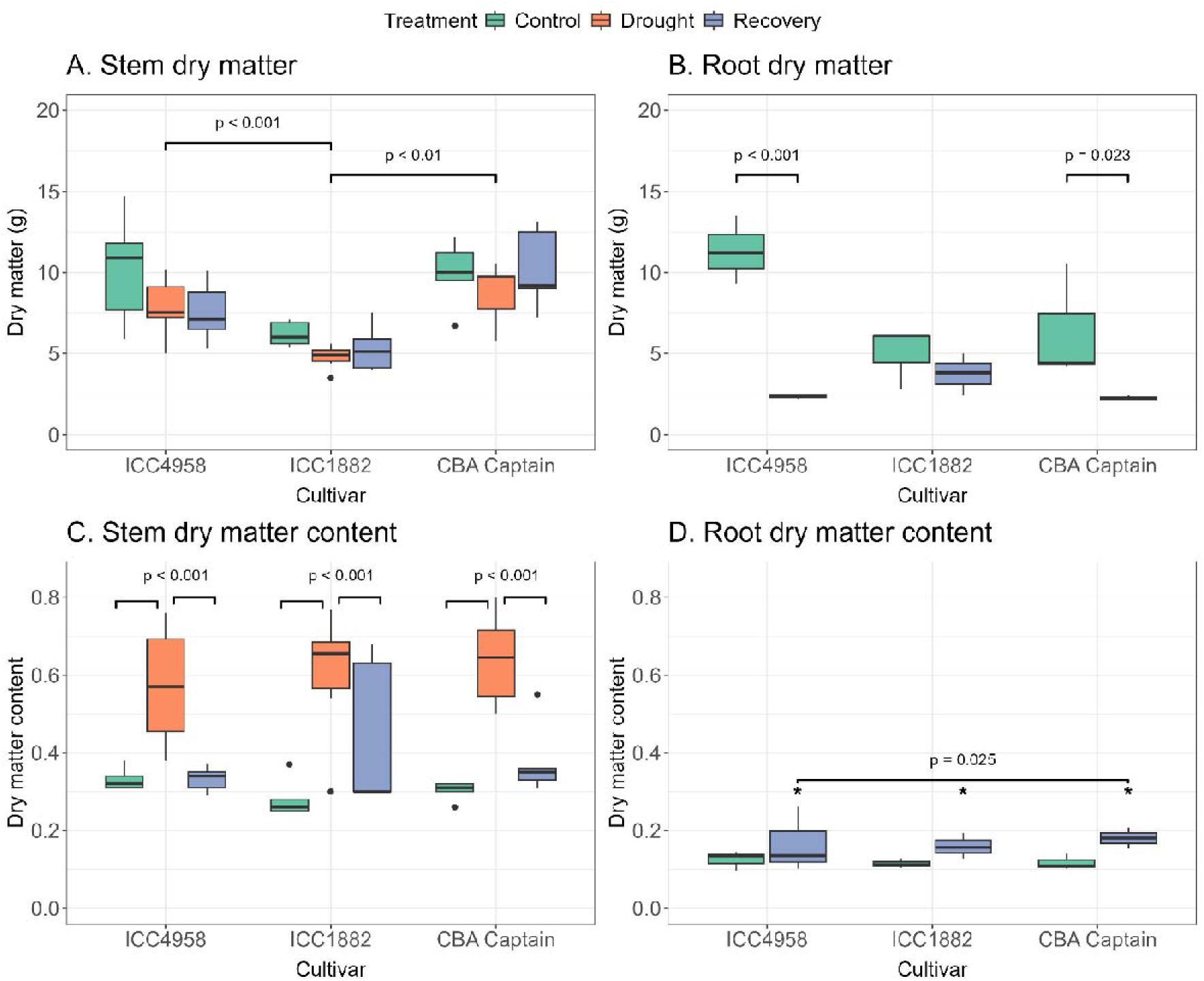
Total stem (**A**) and root (**B**) dry matter per plant (g) measured at the end of the drought experiment as an estimate of harvest biomass (n=3). Stem dry matter was measured for all cultivar and treatment combinations (**A**), with brackets above boxplots indicating results of two factor ANOVA and Tukey’s post-hoc comparisons with p-values indicated where significant (α=0.05). Not enough viable material was present in drought treated replicates to reliably measure root dry matter, however data is shown for Control and Recovery (**B**).

**Figure 4.**
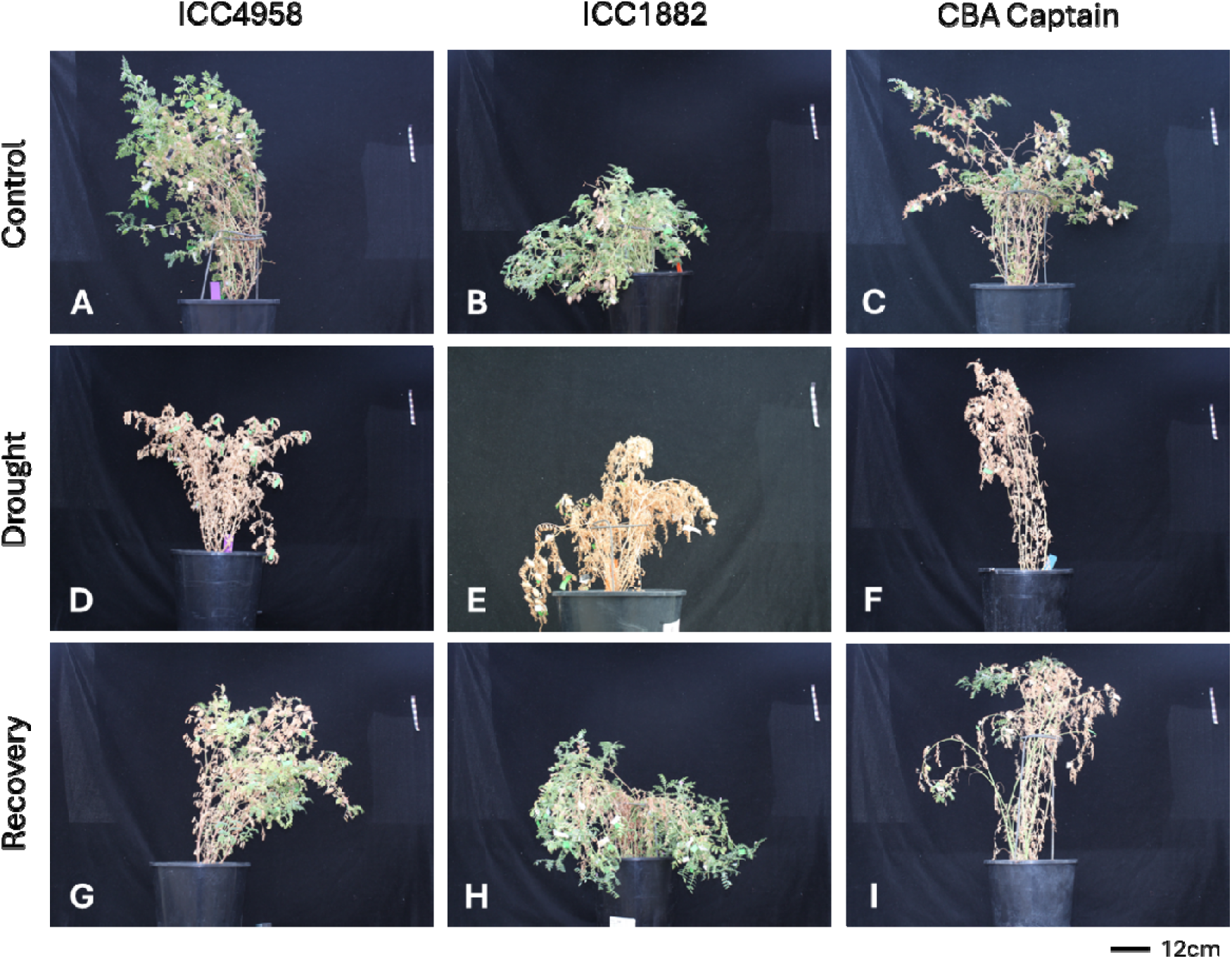
Representative plants from each treatment group and cultivar combination showing their general condition at harvest. The experiment was concluded when CBA Captain Control plants had reached reproductive senescence (**C**). Cultivars are shown in columns and treatment groups in rows. Scale shown in the background (top right) of each image is 12cm as marked at the bottom right.

The impacts of terminal drought treatments were more severe for root biomass than for stems, with the roots of Drought plants being so desiccated and in such a poor condition that it was not possible to isolate them from soil at harvest. The effects of cultivar and treatment were found to interact for root dry matter in remaining treatment groups (p=0.015), however simple main effects analysis within cultivars showed the lower root biomass in Recovery was significant compared with Control for ICC4958 (p<0.001) as well as for CBA Captain (p=0.023) although this was not the case for ICC1882 (**Figure 3B**). In contrast to the findings for stems, the dry matter content of roots was higher in Recovery than Control at harvest (p=0.025) with no significant effect of cultivar (**Figure 3D**). This was also observed visually, where plants in Recovery treatment showed signs of substantial desiccation (**Figure 4G, H, I**) but still had green growth suggesting more viable vegetative tissue than Drought (**Figure 4D, E, F**). Whilst under Control conditions ICC4958 showed a 4.7 fold increase in root dry matter by harvest compared with measurements at peak biomass, the Recovery plants did not undergo this late investment in root biomass with an average of 2.37g almost identical to the 2.40g observed at peak biomass (**Figure 2D and Figure 3B**).

### Dynamic responses to drought stress impact node counts and viability

The effect of treatment on the productivity of podding nodes was highly significant (p<0.01) for both Drought and Recovery which produced fewer grain per plant than Controls regardless of cultivar (**Figure 5B**). Further interrogation of these data by Region however revealed the effects of both cultivar and treatment were varied. In the R1 Control groups CBA Captain produced on average significantly more grain than ICC1882 (p=0.034) whilst the inverse was observed for R2 (p=0.006) in a similar trend to the marginal to significant difference in R3 grain (p=0.058) (**Figure 5B**). These data also suggest the drought stress response of cultivars was not consistent between Regions, with significant differences observed in node productivity at R2 where pods were both formed and filled under treatment. CBA Captain produced significantly fewer grain in R2 under both stress regimes in Drought (p<0.001) and Recovery (p=0.029) groups when compared with ICC4958, as well as lower grain production for Drought than ICC1882 (p=0.009) (**Figure 5B**). Interestingly no significant effects of cultivar were observed in the grain counts for R3, where the Drought groups produced significantly fewer grain than Control for both ICC4958 (p=0.027) and ICC1882 (p<0.001) but not CBA Captain whose entry into senescence was used to signal the end of the trial.

**Figure 5.**
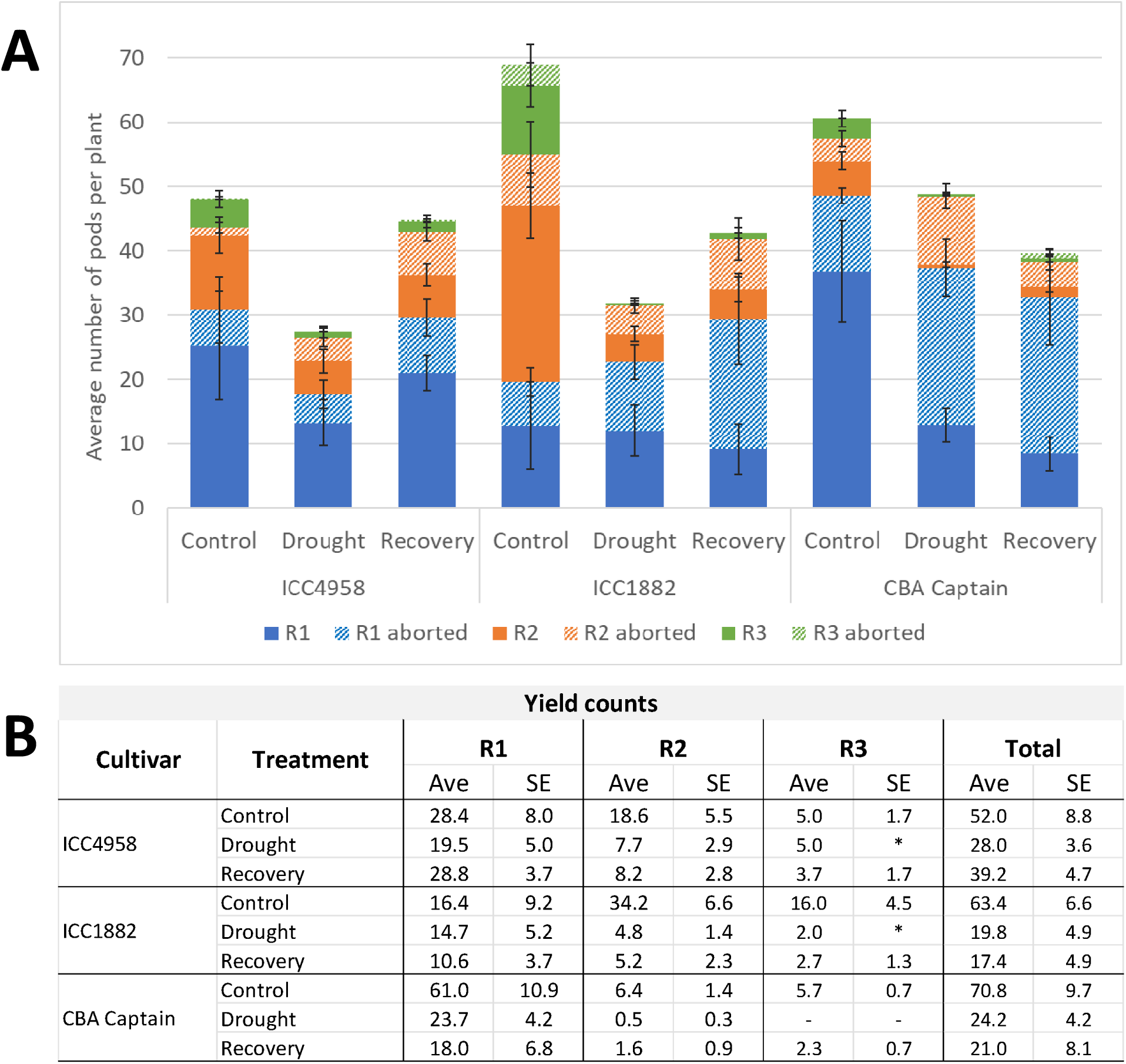
Pods formed per plant by Region for each cultivar and treatment combination (**A**), and the corresponding yield produced (**B**). Viable pods, which contained at least one filled grain, are shown in solid colours whilst corresponding aborted pods in each Region are shown with patterned bars and standard error (SE) indicated (**A**) (n=5/6, +/−SE). The resulting yield by Region is shown as average grain per plant, and SE shown (n=5/6), with the exceptions indicated by asterisk where only a single observation was made in a treatment group (**B**).

Under stress CBA Captain was found to have higher proportions of aborted pods during grain production compared with the other cultivars, indicative of a cultivar effect in drought response. In the absence of drought stress Control treatments showed no difference between cultivars in the proportion aborted pods in any Region. In contrast, in R1 the proportion of aborted pods was significantly higher in CBA Captain than ICC4958 for both Drought (p=0.004) as well as Recovery (p=0.016) (**Figure 5A**). In R2 the proportion of aborted pods in the CBA Captain Drought treatment was significantly higher than either ICC4958 (p=0.011) or ICC1882 (p=0.008) (**Figure 5A**). Surprisingly, when treatment effects were considered by Region there was not found to be a significant difference in the proportion of aborted pods for ICC1882 in either treatment group compared with Control (**Figure 5A**). For both ICC4958 (p=0.043) and CBA Captain (p=0.001) the number of aborted pods in R2 was higher in Drought than Control (**Figure 5A**).

### Continued flowering was observed after rewatering of Recovery plants

For all cultivars Recovery plants were observed to continue to produce green, viable shoot and leaf tissue and, in most cases, new reproductive nodes in the R3 following rewatering (**Figure 6A, B**). Green tissue was observed to emerge from the ends of stems which appeared to be desiccated and senescing under drought stress (**Figure 6A**) as well as from nodes which had previously produced reproductive tissue (**Figure 6B**). The Recovery plants were also able to produce R3 flowers at similar numbers to Controls, with the exception of some ICC1882 individuals which produced more, in contrast to Drought which did not produce any flowers (**Figure 6C**).

**Figure 6.**
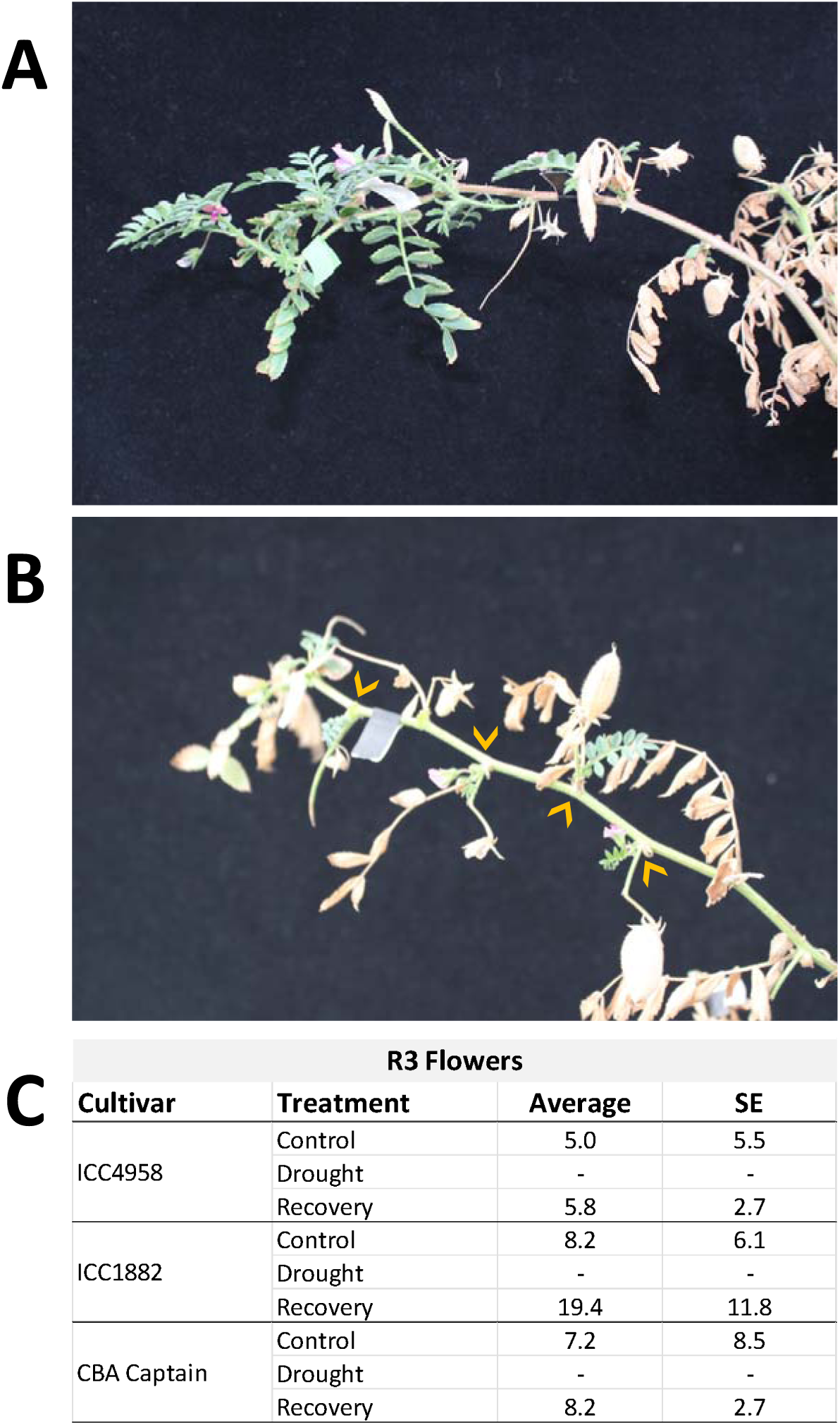
Examples of new growth observed on the desiccated branches of Recovery plants following rewatering, shown for two CBA Captain individuals. New growth including production of flowers in the R3 following rewatering was observed in some cases at the ends of branches (**A**), and in other cases from earlier nodes which had already produced flowers, pods or vegetative growth but which had desiccated under the initial drought period as indicated by yellow arrows (**B**). The average number of R3 flowers is shown for each cultivar and treatment group in the table with SE (n=5/6) (**C**).

### Protein content of grains increases under drought stress

An average of 87-101% of grain weight was accounted for by the combined proximal composition assays which were performed on dried material (**Figure 7A**), not including the moisture content of chickpea grain found to be typically 8.2% in these conditions (Buck et al., 2025). In neither ICC4958 nor ICC1882 were 100 seed weights found to differ between treatments at R1, or indeed other Regions, however CBA Captain produced significantly smaller grain under Drought at R1 (p=0.011) and R2 (p=0.047) as well as in Recovery for R1 (p=0.002) compared with Control (Figure 7B). Across cultivars and treatments the average content of 80% ethanol soluble carbohydrate ranged between 4.5-7.4%, insoluble carbohydrate between 32.6-41.5% and total lipids accounted for 11.7-15.1% of grain weight (**Figure 7A**). More variation was observed in protein and starch content than for other components, accounting for between 13.5-23.1% and 13.5-30.0% of grain weight respectively (**Figure 7A**).

**Figure 7.**
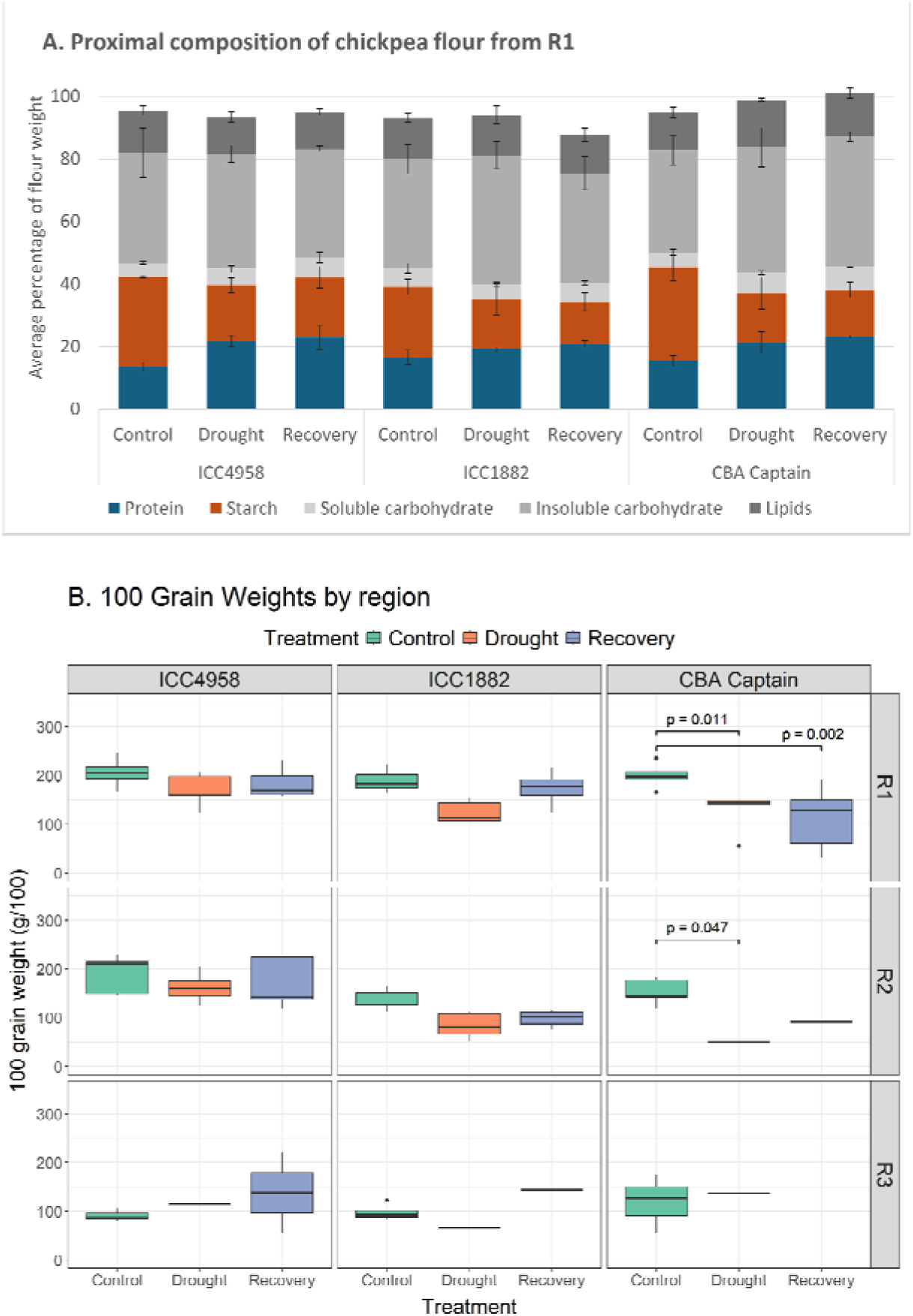
Proximal grain composition of a representative sample of R1 (n=3, +/−SE), which had filled under treatment and reached maturity by the conclusion of the drought experiment (**A**). Grain was assayed for total protein, total starch, 80% ethanol soluble and insoluble carbohydrate, and total lipid with content expressed as average percentage of flour accounted for by each component (w/w) (**A**). The distributions of 100 grain weights (g/100) are shown across all Regions and cultivars (n=5/6), with results of a multifactor ANOVA with simple main effects shown for significant p-values ((α=0.05) (**B**).

The variation observed in protein and starch content of grain was found to be driven by significant treatment effects (MANOVA, p=0.015) whilst there was no significant effect of cultivar in this analysis (**Figure 7A**). Post hoc one-way ANOVA revealed no significant differences in the 80% ethanol soluble or insoluble carbohydrate, or total lipid between treatment groups, however the effect of treatment was highly significant for protein (p=0.002) as well as starch (p=0.001). Tukey’s pairwise comparisons show this effect is driven by a significant, 15.2-60.7% higher protein content in Drought (p=0.016) and 25.7-70.4% higher protein in Recovery (p=0.002) relative to Control, and a corresponding decrease in starch which in Drought (p=0.005) was 28.8-47.2% lower and in Recovery (p=0.003) 33.6-50.2% lower (**Figure 7A**).

## Discussion

The indeterminate flowering habit of chickpea provides an opportunity for the examination of both temporal and developmental impacts of terminal drought on grain production and composition. In this study two broad, dynamic responses to drought stress were observed in reproduction; (1) the consolidation of reproductive nodes by aborting a higher proportion of pods under stress; and (2) the continuation of flowering after alleviation of stress. Significant differences in these responses were observed between the developmental and temporal Regions of flowering, reflecting the importance of timing in even an acute terminal drought stress with rewatering. Poor outcomes for productivity at podding nodes were observed under drought stress regardless of cultivar and were a combination of both altered investment in pod set and filling as well as changes to the quality of grain produced. These changes were consistent across cultivars for composition of grain formed under stress, with a proportionate increase in protein and decrease in starch content observed in R1.

### Cultivars produce different vegetative biomass in response to drought stress

Whilst the broad impacts of drought stress on biomass were consistent across cultivars, most notably in the lower root biomass, there was also variation within cultivars. Differences in phenology confound interpretation of cultivar responses here, where it was necessary to standardise the induction of drought stress and rewatering treatments to entry of one cultivar, CBA Captain, into flowering. For example in Control the ICC1882 plants spend longer in pod set and grain filling so produced most grain in the R2, whereas the two elite cultivars ICC4958 and CBA Captain filled pods earlier, predominantly in R1 (**Figure 5A**). An acute, distinct drought allows characterisation of grain composition with confidence that any changes present were due to treatment, meaning the stress was also defined at slightly offset stages of development for each cultivar.

The growth strategies of cultivars were different under ideal growth conditions, with patterns of investment in biomass reflecting the difference in tolerance or susceptibility to drought stress of ICC4958 and ICC1882. Consistent with previous reports (Garg et al., 2016; Ramamoorthy et al., 2016), during vegetative growth ICC4958 plants accumulated a significantly smaller above ground biomass than both ICC1882 and CBA Captain, measured at peak biomass (**Figure 2A**). Lower specific leaf area during vegetative growth has been associated with higher yields under drought stress in resistant chickpeas including ICC4958, likely due to reduced water loss from the canopy (Ramamoorthy et al., 2016). CBA Captain plants tended to be larger and invest in root biomass earlier, with root mass only increasing 1.27g between peak biomass and harvest compared to the 4.7 fold increase in mass observed in ICC4958. Later investment in root biomass observed in ICC4958 has been documented as a drought avoidance strategy in the variety in field trials (Kashiwagi et al., 2005, 2006; Purushothaman et al., 2016), however this may not extrapolate well to growth in pots. Kashiwagi et al. (2006) also suggest that the ability of resistant cultivars including ICC4958 to establish higher root length density early in vegetative growth allows them to utilise more water at this stage, supporting growth of deeper roots later and resulting in higher yields (Kashiwagi et al., 2006). The earlier peak of root biomass observed in CBA Captain may represent an alternative growth strategy the success of which is contingent on its ability to convert water accessed early to filled grain in the R1 before pressures of terminal drought.

All cultivars experienced severe effects of drought stress in root biomass, although the responses observed in rewatered individuals were varied and are likely caused by differences in favoured growth patterns under ideal conditions. In combination the extensive loss of viable root biomass in Drought plants and the higher DMC of stems is consistent with a severe desiccation and reduction of water supply to aerial tissues in this treatment (**Figure 3C**). That stem DMC was not different between Recovery and Control plants (**Figure 3C**) suggests that aboveground biomass was able to sufficiently rehydrate after the drought treatment. There were however consequences for root biomass, which was found to be significantly lower in Recovery than Control for both ICC4958 and CBA Captain (**Figure 3B**), supporting the conclusion that these plants experienced an intermediate level of drought stress, with rewatering indeed leading to recovery. Recovery plants were able to continue to produce shoot and reproductive tissue at R3 in most cases (**Figure 6**), further evidence that although root biomass was reduced there was still enough water transporting capacity to support new growth. These cultivar differences in accumulation of vegetative biomass, and their consequences for growth during drought stress, impacted grain set and quality by influencing resource availability and the plants capacity for recovery following treatment

### Pod viability is impacted by drought stress regardless of cultivar

Poor outcomes for pod viability and grain filling were driven by stress responses interacting with different growth patterns characteristic of each cultivar, as revealed by the investigation of temporal effects by Region. CBA Captain was found to have the greatest difference in average total grain count between Control and Drought, this was not found to be a statistically significant (**Figure 5B**). The average total grain production in ICC4958 Drought and Recovery were slightly higher than for ICC1882 and CBA Captain but this difference was not pronounced enough to be significant (**Figure 5B**). Although not significant in this study, it is possible that differences in grain production could represent more subtle cultivar effects that would be captured by larger sample sizes. There were significant effects of Region however that were most evident in CBA Captain which produced significantly fewer R2 grain, from pods that were both set and filled under stress, in Drought and Recovery compared with the other cultivars (**Figure 5B**). Under well-watered conditions CBA Captain tended to produce the majority of its grain earlier than other cultivars, as such the acute drought impacted its most highly productive stage, although some R1 pods had been filled before the imposition of stress (**Figure 5**). In ICC4958 R2 is more productive in well watered conditions than for CBA Captain, contributing a higher fraction of total grain filled that reflects a more steady rate of grain production across flowering (**Figure 5**).

Pod abortion was a strategy deployed by both CBA Captain and ICC4958 under Drought conditions, with significantly higher proportions of aborted pods in R2 than Control (**Figure 5A**) suggesting that consolidation of reproductive nodes to those already filling was initiated as an adaptive response to stress. This observation is consistent with several previous studies which report increases in the proportions of aborted pods in terminal drought where stress is applied early in podding, leading to reduced grain filling (Fang et al., 2010; Hossein Behboudian et al., 2001; Leport et al., 2006; Pang et al., 2017). The impact of pod abortion was more pronounced for CBA Captain which aborted higher proportions of pods in both R1 and R2 than ICC4958 (**Figure 5A**), raising the possibility that an induction of drought slightly earlier than in this experiment would have even more devastating impacts on grain filling for this cultivar. These data suggest that ICC4958 is less likely to abandon pods that have already set than CBA Captain, as only the R2 which were initiating under stress showed significantly higher proportions of aborted pods and not the R1 which would have a higher proportion that were already committed to filling (**Figure 5**).

There is also a more apparent reduction in the number of nodes which progressed to pod set at all, whether viable or aborted, for ICC4958 Drought compared with Control suggesting that this cultivar is less likely to invest in producing more reproductive nodes under stress than CBA Captain (**Figure 5**). This is interesting in the context of reports that pollen viability is reduced under terminal drought which inhibits germination and leads to abnormal pollen tube development (Fang et al., 2010; Pang et al., 2017). Further investigations would be needed to resolve if pollen viability was impacted under these conditions, or whether this reflects a more conservative strategy for ICC4958 in committing to further development of new reproductive nodes than the more reactive response observed in CBA Captain. In contrast ICC1882 was not found to have higher proportions of aborted pods in any Region under either stress treatment, indicating this was not a strategy deployed in response to stress. Although consolidation of resources to committed reproductive nodes may help secure grain already filling, this strategy would not allow plants to quickly resume production at the alleviation of stress without forming new reproductive nodes, consistent with the observation of new growth at R3 in most Recovery plants (**Figure 6**).

### Terminal drought stress is associated with changes in grain composition

In addition to reduced ability to fill grain under terminal drought stress the macronutrient composition of grain that is produced changes, with an increase in total protein and a decrease in starch (**Figure 7A**). Changes to grain composition are driven by effects of drought stress in both treatments and not by cultivar, suggesting these findings are likely to apply more broadly to other varieties of chickpea. Although a reduction of 100 grain weight was observed under Drought and Recovery for CBA Captain in R1 (**Figure 7B**), the absence of a cultivar effect on grain components suggests this did not influence the proportion of each in grain. Proximal composition analysis was performed only on R1 grain, because this material had time to complete grain filling prior to harvest to avoid introducing variability with sampling different developmental stages (**Figure 1**). The R1 grain were filled under stress treatments, parallel to changes that were observed in development and biomass allocation, so represent grain specific consequences of broader stress responses.

The converse relationship observed in this study between protein and starch in grain under stress differs from previous reports in the limited number of studies which have considered all the major grain components under drought. These studies applied drought treatments at later stages of pod filling than in experiments presented here, introducing the possibility that temporal effects determined different outcomes. In addition, this study is the first to consider the developmental stage of grain in relation to the applied stress. However, comparisons between studies are fraught due to differences in application of stress, in particular the use of a combination of field and cabinet conditions by Awasthi (2024), and different total proportions of flour accounted for in composition analyses. This highlights the need for further studies to investigate the temporal effects of drought on grain composition in consistent, controlled environments.

Evidence from field trials with Kabuli chickpeas is consistent with findings presented here, which suggests availability of sufficient water prior to flowering is a key to determining grain protein, even if drought stress is experienced during filling (Benali et al., 2023; Samineni et al., 2022; Varol et al., 2020). Applying different irrigation strategies Varol et al. (2020) report the lowest grain protein content in rainfed plants with the greatest increase in groups which received irrigation immediately preceding flowering. The total starch in grain was found to be positively correlated to the amount of water supplied during reproduction, highest in groups that received water at the beginning of flowering (Varol et al., 2020). In recent investigations of heat stress and combined heat and drought stress Benali et al. (2023) observed that grain protein increased to the greatest extent in plants which received water until flowering before induction of combined stress. Authors suggest that increased expression of stress related proteins may be causing observed increases in protein (Benali et al., 2023), which cannot be excluded although storage proteins typically account for the majority of this component of grain. These reports of higher protein content produced by treatments which received sufficient water into flowering are consistent with the observation that Drought and Recovery plants showed no significant difference in any grain component, despite the resumption of watering during grain filling in the latter (**Figure 7A**).

In combination these data suggest the most likely driver of changes to grain composition under terminal drought is the temporal effects on starch deposition, and that differences in protein are relative to alterations in carbohydrate metabolism. There is increasing evidence that for chickpea, as in other plants, the photosynthetic rate decreases under drought stress and that this has implications for the supply and utilisation of sucrose to developing grain (Awasthi et al., 2014, 2024; Nayyar et al., 2006). In two very similar experiments, the photosynthetic capacity of chickpea under drought stress during grain filling is reduced and consequently the sucrose content of leaf and grain decreases, whilst soluble sugars increased in grain (Awasthi et al., 2014, 2024). Key aspects of carbohydrate metabolism are altered including reduced activity of enzymes crucial to utilising sucrose for synthesis and deposition of starch in the grain including sucrose synthase, acid invertases and starch synthase, as well as turnover of starch including β-amylase and starch phosphorylase (Awasthi et al., 2014, 2024; Nayyar et al., 2006). This suggests that the availability of photoassimilates, mobilisation of sucrose and conversion into carbohydrates in the grain is compromised under drought stress in chickpea, resulting in reduced starch deposition (Awasthi et al., 2014, 2024; Nayyar et al., 2006). Changes under drought to carbohydrate metabolism more broadly have been documented in recent multi-omics studies of roots in ICC4958 and ICC1882 (Kudapa et al., 2024; Singh et al., 2023) as well as vegetative growth in other cultivars (Farooq et al., 2018) and suggested to be important in mechanisms of tolerance. That starch was found to decrease in the grain of Drought and Recovery groups in all cultivars (**Figure 7A**) is consistent with these previous reports that carbohydrate metabolism is altered under stress and has consequences for grain filling. It is interesting that this effect occurs regardless of rewatering in the Recovery plants, which suggests that even a short disruption early in podding has substantial effects on the filling of grain. The disruption of carbohydrate production leading to lower deposition of starch during filling may have resulted in proportionately more protein in grain, a component which was less impacted by drought stress as water supply prior to flowering was sufficient in all treatments.

## Conclusions

The productivity of reproductive nodes under terminal drought was found to be strongly influenced by the timing of this stress relative to development, as governed by cultivar specific patterns of growth. This was most evident in the decreased rates of reproductive node initiation and increased pod abortion in Regions forming under stress for both CBA Captain and the drought tolerant ICC4958. These data demonstrate the importance in examining both temporal and developmental effects of drought stress to understanding reproductive responses in chickpea. Regardless of cultivar the composition of grain resulting from nodes that developed under stress show increased protein and decreased starch as a proportion of total grain weight, consequently impacting their nutritional content. Whilst available evidence suggests this likely reflects altered carbon metabolism resulting from reduction in available photoassimilates, there is not enough currently enough known about the mechanisms of grain filling in chickpea to fully interpret these changes in grain composition.

## Acknowledgments

Authors would like to acknowledge discussions with Dr Alex Whan in the selection of data analyses. Paul Hopgood was funded by the Grains Research and Development Corporation as partners in the CSIRO Summer Scholar program.

